# A late Permian ichthyofauna from the Zechstein Basin, Lithuania-Latvia Region

**DOI:** 10.1101/554998

**Authors:** Darja Dankina-Beyer, Andrej Spiridonov, Ģirts Stinkulis, Esther Manzanares, Sigitas Radzevičius

## Abstract

The late Permian is a transformative time, which ended in one of the most significant extinction events in Earth’s history. Fish assemblages are a major component of marine foods webs. The macroevolution and biogeographic patterns of late Permian fish are currently insufficiently known. In this contribution, the late Permian fish fauna from Kūmas quarry (southern Latvia) is described for the first time. As a result, the studied late Permian Latvian assemblage consisted of isolated chondrichthyan teeth of *Helodus* sp., ?*Acrodus* sp., ?*Omanoselache* sp. and euselachian type dermal denticles as well as many osteichthyan scales of the Haplolepidae and Elonichthydae; numerous teeth of *Palaeoniscus*, rare teeth findings of ?*Platysomus* sp. and many indeterminate microremains. This ichthyofaunal assemblage is very similar to the contemporaneous Lopingian complex of the carbonate formation from the Karpėnai quarry (northern Lithuania), despite the fact that Kūmas samples include higher diversity and abundance in fossil remains. The differences in abundance of microremains could possibly be explained by a fresh water influx in the northeastern Zechstein Basin margin, which probably reduced the salinity of the sea water. The new data enable a better understanding of the poorly known late Permian fish diversity from the Lithuania-Latvia Region.

## Introduction

The late Permian is currently one of the most studied time periods in the whole Phanerozoic, since it is evidenced by two phases of the end-Permian mass extinction, which significantly affected long term macroevolution of biota [1]. Despite an overall low level of detailed taxonomic palaeoichthyofaunal analysis there is an increased interest on the late Permian fossil fish assemblages, which is mostly focused on the isolated chondrichthyan and osteichthyan remains (dermal denticles, scales, fins, vertebral centra, teeth) demonstrating their usefulness for palaeoenvironmental and palaeoecological reconstructions of complex marine vertebrate ecosystems from the time period immediately preceding Permian-Triassic mass extinction [2–3].

Osteichthyans and chondrichthyans represent the dominant fish classes among marine and freshwater vertebrates since the Carboniferous period [3–4] and are relatively common in the late Permian fossil records. Fish remains occur widely around Pangea, including current areas of Australia [5], South Africa [6–8], Brazil [9–10], China [11–13], Japan [14], Russia [15–17], Iran [18], India [19] and Greece [20]. Moreover, various fish assemblages were widely spread in the Zechstein Basin, based on the fossil findings from Germany [21], England [22], East Greenland [23], Poland [24] and Lithuania [25].

The Zechstein Basin is characterized by seven evaporite cycles each of which starts with clastic sediments followed by carbonates, sulfates, chlorides and finally potash salt [26]. However, evaporite cycles are not complete in the Lithuania-Latvia Region and are represented by only carbonates which are lithostratigraphically related to a part of the Zechstein cycle Nº1 [27].

The current study represents the first record of a fish assemblage from Sātiņi, Kūmas, and Alši Formations in Kūmas quarry, southern Latvia – the north-easternmost part of the Zechstein Basin, never before studied on the palaeoichthyofaunal grounds. The isolated remains of Palaeoniscidae, Platysomidae, Haplolepidae, Elonichthydae and Euselachii, Helodontidae, Acrodontidae were revealed from the carbonate deposits of Zechstein Limestone (Ca1) formed by the first cycle Werra (Z1) [27–28].

The main goal of this paper is taxonomic designations of the fish assemblage of the Zechstein Basin. We also present computed tomographic and histologic models of the internal vascular structures of these isolated microvertebrate fossils. The purpose of these 3D and histological modelling is to provide additional data for future taxonomic identifications. This material was also compared to the geographically proximate Karpėnai quarry in northern Lithuania [25].

## Geological setting and stratigraphy

The Southern Permian Basin extends over a distance of 1700 km from England across the Southern North Sea through Northern Germany into Poland and the Baltic States [29]. The most northeastern nearshore deposits of the studied Basin are located Lithuania and Latvia.

In northern Lithuania, close to Naujoji Akmenė city, Karpėnai quarry is a functioning cement quarry (Fig 1). The late Permian deposits in the quarry consist of three members of Naujoji Akmenė Formation which are represented by different types of carbonate sediments from Zechstein Limestone (Ca1). The Lower Member is characterized by a 2 m thick micritic limestone, the Middle Member consists predominantly of the 8 m thick proximal tempestites and the Upper Member is a 4 m thick limestone with coated grains, where in the highest part of the unit consist of dolomitic limestone [25].

**Fig 1.**
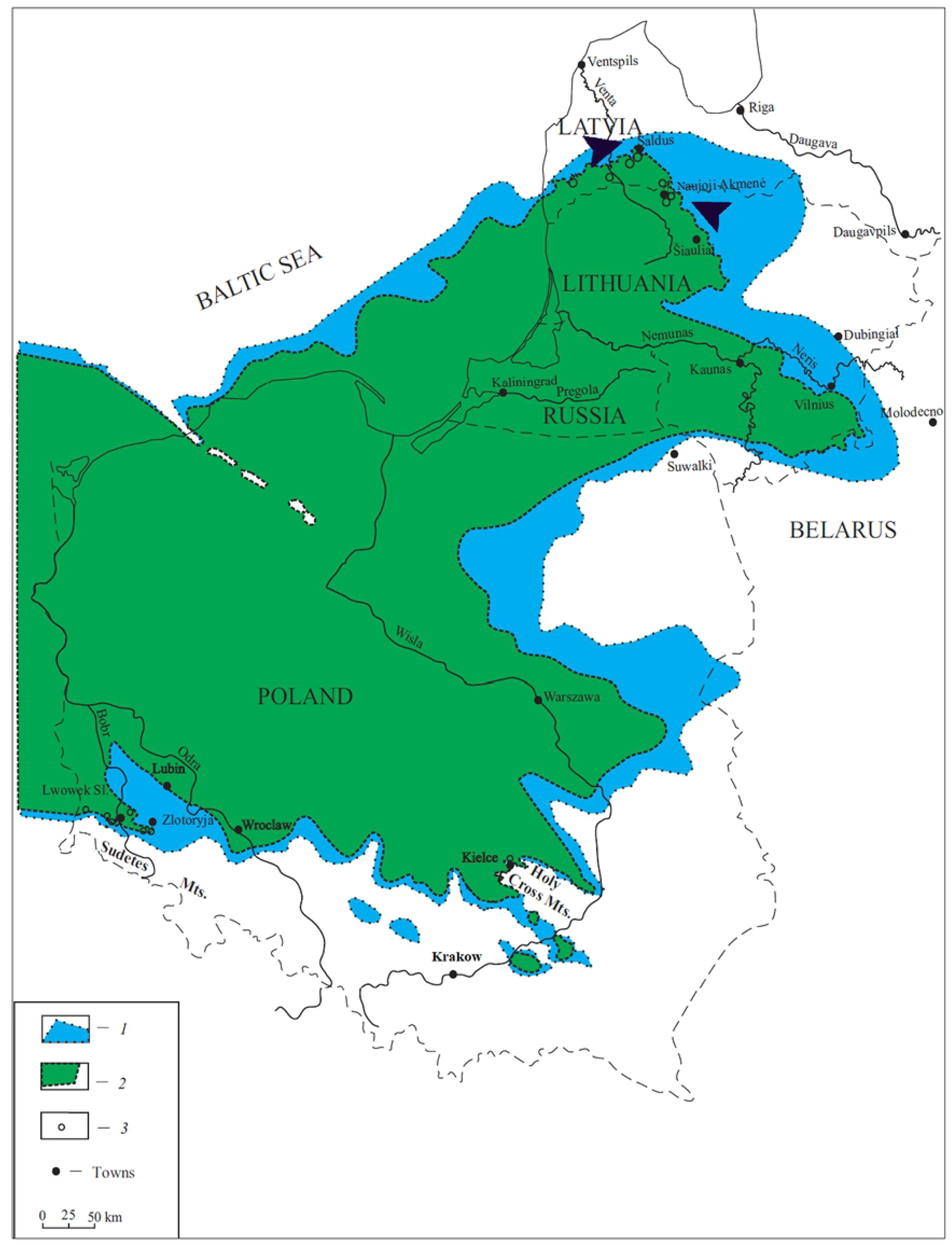
Geographical map of the eastern part of the Zechstein Basin (modified by Raczyński and Biernacka, 2014). (1) original; (2) current distribution of Zechstein sediments in Poland (PL), Lithuania (LT) and Latvia (LV). (3) Zechstein outcrops. (Black arrows), showing two studied locations of the Karpėnai and Kūmas quarries.

Kūmas quarry is situated in the south-western part of the Latvia (Fig 1). Limestones of the Permian, Werra Series, Naujoji Akmenė Formation [27] are represented here by three members of the Naujoji Akmenė Formation: Sātiņi, Kūmas, and Alši ones [30–31]. Sandy limestones with bivalve fossils of the Sātiņi Member, mostly less than 2 m thick, are not available in the main territory of the quarry and are exposed only in water collecting ponds represented the lowermost part of Naujoji Akmenė Formation. They directly cover the lower Mississippian (Tournaisian), sandstones. The Sātiņi Member is covered by the earthy limestones with nodules and interlayers of porcelain-like limestones of the Kūmas Member. This section of the quarry reaches a thickness of 10-14 m. The fossiliferous limestones of the Alši Member are 5-7 m thick in the territory of the Kūmas quarry. The total thickness of the Naujoji Akmenė Formation in this quarry varies from 15 to 22 m [30]. The Permian limestones in the Kūmas quarry are covered by the Quaternary, mainly till deposits, reaching thickness of 10-20 m (Fig 2).

**Fig 2.**
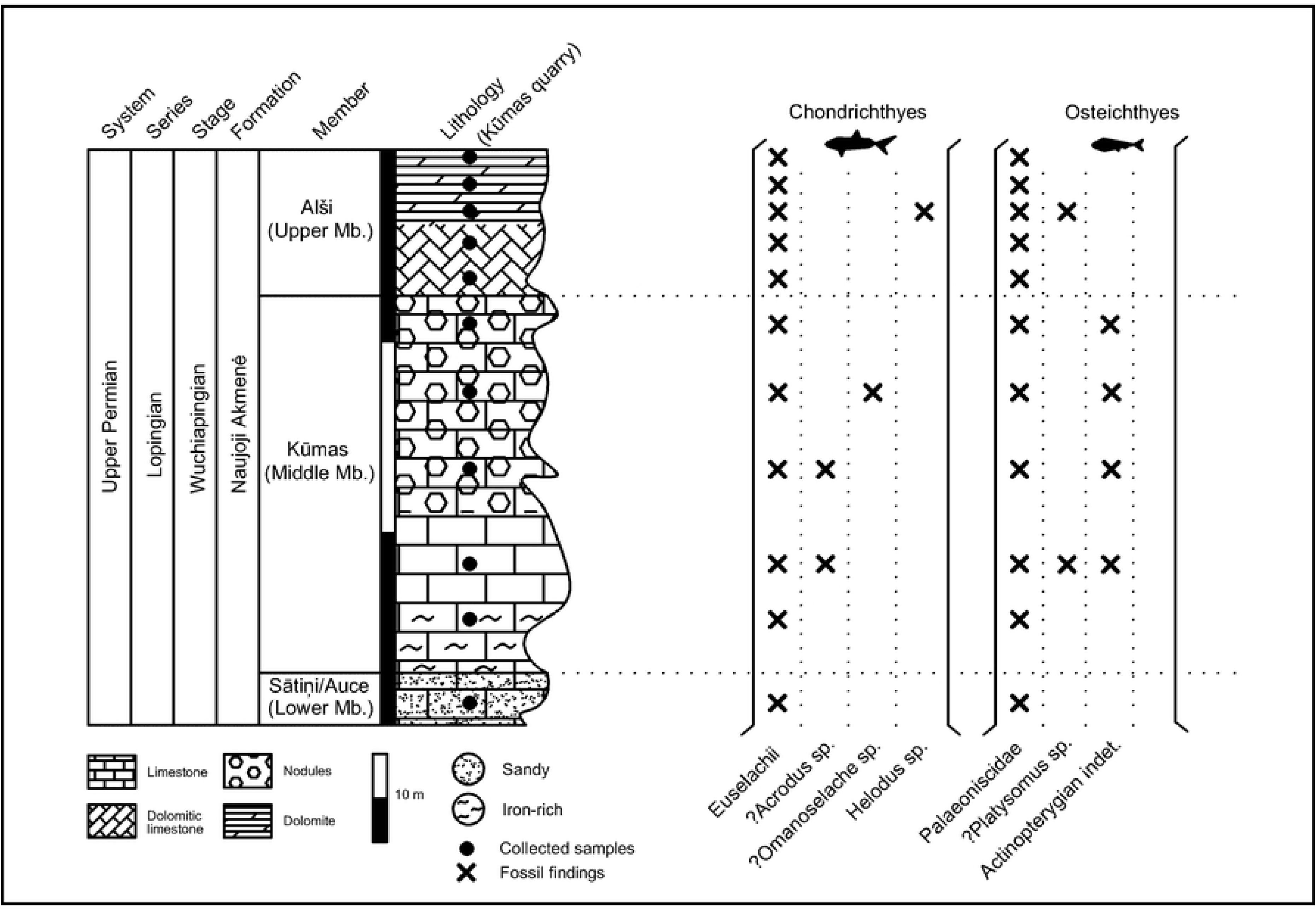
Stratigraphical profile of the Kūmas quarry (southern Latvia) with indication of the late Permian fish assemblage, and stratigraphic repartition of the chondrichthyan and osteichthyan taxa based on isolated teeth, dermal denticles and scales.

The biostratigraphical correlation between earlier described quarries in the Lithuania-Latvia Region shows us the same pattern of the first and last occurrences of cartilaginous and bony fish isolated fossils, which could point to similar palaeoecological, taphonomic and sedimentological mechanisms responsible for their accumulation (Fig 2). In the Lower Member of Naujoji Akmenė Formation actinopterygian scales, teeth and chondrichthyan dermal denticles were found based on the analysis of both outcrops. The Middle Member is represented by common findings of euselachian type dermal denticles, actinopterygian teeth and rare *Helodus* sp. teeth, while *Palaeoniscus* scales are nearly absent in both quarries. Although, there are few ?*Acrodus* sp. teeth microremians in the Kūmas quarry. Actinopterygian teeth and dermal denticles were found in the Upper Member in both quarries whereas *Palaeoniscus* scales are presently revealed only in the Kūmas quarry (Table 1, Fig 2). According to the statistical count data, the higher variety and number of microremains are present in Kūmas quarry.

**Table 1.** Abundance of palaeoichthyofaunal microremains from the Kūmas quarry, compared with the sample locations in the outcrop (from the ground level), weight, weight estimates of dissolution residues.

| Nr. | Specimen | Height | Weight | Residue | Actinopterygian |  | Chondrichthyan |  |
| --- | --- | --- | --- | --- | --- | --- | --- | --- |
|  |  | (m) | (kg) | (g) | Tooth | Scale | Tooth | D. denticle |
| 1 | KUM-11 | 14.0-15.0 | 18.4 | 44.02 | 215 | 1 | 0 | 64 |
| 2 | KUM-10 | 13.0-14.0 | 15.5 | 32.45 | 17 | 0 | 0 | 2 |
| 3 | KUM-9 | 12.5-13.0 | 28.3 | 93.24 | 23 | 1 | 1 | 1 |
| 4 | KUM-8 | 12.0-12.5 | 16.5 | 170.01 | 4 | 1 | 0 | 1 |
| 5 | KUM-7 | 11.5-12.0 | 24.1 | 91.86 | 6 | 1 | 0 | 2 |
| 6 | KUM-6 | 10.5-11.5 | 23.4 | 77.52 | 16 | 1 | 0 | 2 |
| 7 | KUM-5 | 7.5-10.5 | 29.3 | 57.25 | 31 | 0 | 1 | 1 |
| 8 | KUM-4 | 5.0-7.5 | 18.5 | 19.08 | 220 | 35 | 3 | 73 |
| 9 | KUM-3 | 2.5-5.0 | 24.7 | 8.02 | 939 | 109 | 13 | 79 |
| 10 | KUM-2 | 0.5-2.5 | 10.6 | 34.29 | 209 | 57 | 0 | 14 |
| 11 | KUM-1 | 0.0-0.5 | 10.3 | 90.44 | 13 | 5 | 0 | 2 |
| **Total:** |  | 15.00 | 219.6 | 718.18 | 1693 | 211 | 18 | 241 |

In the studied area, we found many unidentified vertebrate fossils (e.g. fragment of a conodont element, isolated fish vertebral centra, otoliths and tooth plate), invertebrate and plant fossils (e.g. spores and pollens), which need additional detailed taxonomic analysis.

## Material and methods

Material used in this study was excavated in the November 2016 in the Kūmas quarry (56°34’56”N; 22°35’82” E), in southern Latvia. The material presented here is compared to earlier published report on the palaeoichthyofauna of the Karpėnai quarry, norther Lithuania [25]. The geographical distance between these locations is ∼70 km. The total weight of the analysed samples from Karpėnai quarry reached ∼214 kg and from Kūmas quarry - ∼220 kg. These samples yielded ∼2,400 isolated fish microremains from both quarries (Table 1).

The samples were chemically prepared using the optimal acetate 10 *per cent* buffered acetic acid technique for extracting phosphatic fossils [32] in the Micropalaeontological Laboratory in Institute of Geosciences at Vilnius University (Vilnius, Lithuania). Fish microremains were picked from residue samples onto microslides. Some of the microremains were photographed using a HITACHI S4800-FEG scanning electron microscopy (SEM) at the University of Valencia (Valencia, Spain) and a FEI Quanta 250 SEM model in the Nature Research Centre (Vilnius, Lithuania).

For histological analysis specimens were ground and polished then embedded in a transparent polyester resin at 120°C for 30-45 minutes prior to polishing with carborundum grain (1200 µm) and water until the desired part of the fossil was reached. Afterwards, the longitudinal and transversal sections were etched for 5-10 s in 10 *per cent* HCl acid. Each sample was re-polished and etched as many times as necessary to elucidate the enameloid microstructure [33]. The ground sections were coated with gold-palladium prior being photographed using a Hitachi S4800-FEG SEM at the Microscope Service at the University of Valencia.

Two fossils from Karpėnai quarry were scanned using Synchrotron Light Source from Paul Scherrer Institute (PSI) in Zurich, Switzerland for internal morphology [34–35]. Scanning of the entire *Helodus* sp. tooth [VU-ICH-KAR-012] produced 930 2D sections with 1.625 μm voxel size units; and the euselachian dermal denticle [VU-ICH-KAR-023] produced 2,270 2D sections with 0.325 μm voxel size units. These sections were processed with Avizo 8.1 software [36] in order to generate and reconstruct 3D models at the University of Valencia (Valencia, Spain).

All material presented here (numbered VU-ICH-KUM-001 to VU-ICH-KUM-059) is reposited in the Geological Museum of the Institute of Geosciences at Vilnius University.

## Systematic palaeontology

Class CHONDRICHTHYES Huxley, 1880 [37]

Subclass ELASMOBRANCHII Bonaparte, 1838 [38]

Order HYBODONTIFORMES Patterson, 1966 [39]

Family ACRODONTIDAE Casier, 1959 [40]

Genus *ACRODUS* Agassiz, 1838 [41]

**Type Species**—*Acrodus gaillardoti* Agassiz, 1839 [41]

*?ACRODUS* sp. (Fig 3A-B)

**Fig. 3.**
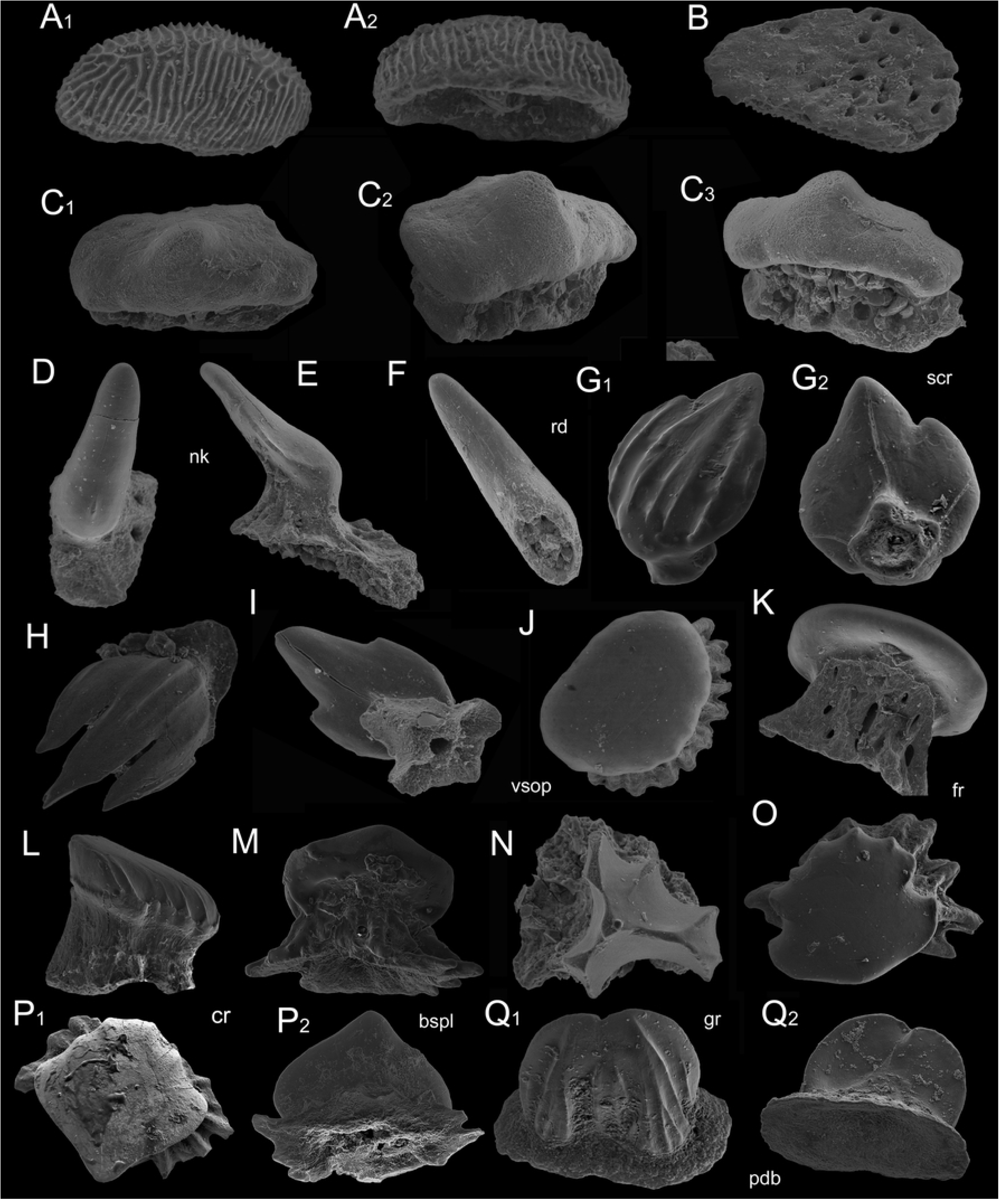
SEM images of the late Permian chondrichthyan dermal denticles, *?Acrodus* sp. and ?*Omanoselache* sp. teeth from Kūmas quarry. (A) *?Acrodus* sp. tooth, VU-ICH-KUM-001 from KUM-3 sample. (A_1_) tooth in occlusal view; (A_2_) in oblique lateral view. (B) VU-ICH-KUM-002 from KUM-3 sample in basal view. (C) *?Omanoselache* sp. tooth, VU-ICH-KUM-003 from KUM-3 sample. (C_1_) tooth in occlusal view; (C_2_) in oblique occlusal view; (C_3_) in lateral view. (D–F), Morphotype 1. (D) VU-ICH-KUM-004 in occlusal view, from KUM-4 sample. (E) VU-ICH-KUM-005 in lateral view, from KUM-3 sample. (F) VU-ICH-KUM-006 in subcrown view, from KUM-3 sample. (G–I) Morphotype 2. (G) VU-ICH-KUM-007, from KUM-2 sample, (G_1_) in occlusal view, (G_2_) in basal view. (H) VU-ICH-KUM-008 in occlusal view, from KUM-10 sample. (I) VU-ICH-KUM-09 in basal view, from KUM-10 sample. (J–M) Morphotype 3. (J) VU-ICH-KUM-010 in occlusal view, from KUM-3 sample. (K) VU-ICH-KUM-011 in lateral view, from KUM-3 sample. (L) VU-ICH-KUM-012 in lateral view, from KUM-11 sample. (M) VU-ICH-KUM-013 in lateral view, from KUM-2 sample. (N) Morphotype 4. (N) VU-ICH-KUM-014 in occlusal view, from KUM-10 sample. (O–P) Morphotype 5. (O) VU-ICH-KUM-015 in occlusal view, from KUM-2 sample. (P) VU-ICH-KUM-016, from KUM-2 sample. (P_1_) in occlusal view; (P_2_) in oblique basal view. (Q) Morphotype 6. (Q) VU-ICH-KUM-017, from KUM-2 sample. (Q_1_) in occlusal view; (Q_2_) in oblique basal view. Abbreviations: (bspl) basal plate; (cr) crown; (fr) foramina; (gr) grooves; (nk) neck; (pdb) pedicle base; (rd) ridges; (scr) subcrown; (vsop) vascular opening. Scale bars equal 0.1 mm respectively.

## Material

16 isolated teeth, only 2 teeth with preserved roots, were found in Kūmas quarry, southern Latvia. This genus represents and described here by a tooth without root [VU-ICH-KUM-001] and tooth with preserved roots [VU-ICH-KUM-002].

## Description

Teeth crown outline has convex ellipse shape, with well-developed sculptured surface made out of the coarse-anastomosing ridges originating from the breadthways crest to mesial juncture in the centre (Fig 3A_1_). It reaches 0.2 mm in length and 0.05 mm in width. The flat basal plate repeats crown outline, preserved roots have numerous canal openings in proximal view (Fig 3B), and however, most common roots are absent (Fig 3A_2_).

## Remarks

Morphological similarities of the crown can be detected with *Acrodus kalasinensis* tooth from the Late Jurassic-Early Cretaceous of northeastern Thailand [42]. Also, similar teeth but evidently bigger were found and described as *Acrodus spitzbergensis* teeth from the Lower Triassic of Spitsbergen [43].

Family INCERTAE SEDIS

Genus *OMANOSELACHE* Koot et al., 2013 [44]

**Type Species**—*Omanoselache hendersoni* Koot et al., 2013 [44]

*?OMANOSELACHE* sp. (Fig 3C)

## Material

Isolated tooth [VU-ICH-KUM-003] from Kūmas quarry, southern Latvia.

## Description

The tooth crown outline is robust, elongate, generally smooth, has flattened appearance in lingual-lateral view and more convex in labial-lateral view. The crown surface has clear asymmetry (Fig 3C_1_) and reaches 0.7 mm mesio-distally, 0.3 mm in labio-lingually, about 0.35-0.4 mm high. The central convex groove has transverse direction with a single blunt, small peg located in the labial margin (Fig 3C_2_). The neck is massive, broad, with some foramina canal openings filled by ‘rubbish’ (Fig 3C_3_). The basal plate is ellipse shape, reaches 0.7 mm in length and 0.3 mm in width.

## Remarks

Similar morphological features of the 3 times bigger tooth were found from the Knuff Formation of Middle Permian in Oman [44]. However, the quantity of the compared material is generally lacked.

Cohort EUSELACHII Hay, 1902 [45]

EUSELACHII indet.

(Fig 3D-Q)

## Material

241 isolated chondrichthyan dermal denticles were found in Kūmas quarry, southern Latvia. These were classified into five morphotypes based on their morphological features and represented here in the description by 14 chosen dermal denticles [VU-KUM-ICH-004 – VU-KUM-ICH-017].

## Description

Well-preserved fossils are identified as euselachian type dermal denticles based on the resembling material from the Upper Jurassic of Tanzania [46], from Middle and Late Triassic of northeastern British Columbia, Canada [47], from the Middle Permian of Apache Mountains, West Texas (USA) [48] and divided into several morphotypes based on their morphological difference between fossil crowns, necks and pedicle base outlines.

### Morphotype 1

This is the most scarce of all morphotypes (3 microremains) in the Kūmas quarry (Fig 3D-F). The crown is blunt horn-like shaped, elongate, slightly depressed in lateral view and smooth: with a high neck and wide base (Fig 3D-E). The crown and base lengths are similar – about 0.5 – 0.6 mm - but direction of the growth is opposite (Fig 3E). The crown is slightly narrower than the neck - 0.2 mm. The pedicle base is sometimes absent (Fig 3F) or, in the case that is present here, has some foramina of vascular system which are usually filled up by sediment.

### Morphotype 2

This is one of the most common type of dermal denticle in the Kūmas quarry (Fig 3G-I). A trident or nearly trident crown with a low, slender and narrow neck, hidden under the crown in dorsal view (Fig 3G_1_,H). The pedicle base has a rhomboid flat outline and one roundish canal opening in proximal view (Fig 3G_2_,I). The exterior of crown is sculptured with some (usually five) gentle convex ridges and furrows originating from the longitudinal crest and reaches 0.3 – 0.4 mm length. The crown sits horizontally or slightly obliquely up on the neck. The subcrown is smooth or has one mesial and two lateral lines (keels).

### Morphotype 3

Roundish or semi-roundish dermal denticle with slightly elongate blunt cusp in posterior margin of the crown (Fig 3J-M). The crown surface is thick, without ornamentation or covered by some short convex ridges and furrows in the anterior side (Fig 3J,L). The crown sits horizontally or slightly obliquely up on the neck (Fig 3L) and reaches about 0.4 – 0.8 mm in diameter. The neck is relatively very high, massive and has numerous foramina of vascular system (Fig 3K-M). The wide pedicle base outline is curved, multipetaloid shaped with concave canal openings in proximal view (Fig 3M). Similar roundish scale was interpreted as a Hybodont/synechodontiform? scale from the Lower Triassic of Oman by Koot et al. (2015, fig. 11A-B). Also, an elongate semi-roundish scale (similar to the ones described in this work) was found and assigned to Hybodontidae gen. et sp. indet. from the Triassic of Spitzbergen [49].

### Morphotype 4

Very rare dermal denticles looks as ‘tripod boomerang’, with three main lateral cusps, two of which separated by two grooves more (Fig 3 N). The crown sits strongly horizontally on the slender neck. The flat pedicle base is poorly-preserved, has unclear shape. Such dermal denticle reach 0.2 – 0.3 mm in length.

### Morphotype 5

Very common euselachian type of dermal denticles in the Kūmas quarry (Fig 3O,P_1_-P_2_). The well-preserved drop-like crown margin outline, smooth, with some short ridges in anterior side which sits evident obliquely up on the wide neck (Fig 3P_1_). The crown length of the mesial platform (positioned at the centre of upper crown surface and runs anterior to posterior) reaches 0.5 mm. The neck and crown width almost the same, about 0.3 – 0.4 mm. The pedicle base outline is curved, multipetaloid shaped with concave canal openings in proximal view. The subcrown is smooth, without any ornament (Fig 3P_2_). Similar dermal denticle crown surface with features was assigned as hybodontiform scale from the Middle Triassic of Spain [33]. Less common crown surface outline is smooth, flat, ‘anchor-like’ shaped with roundish posterior margin, while anterior has 6 short convex ridges and two more evident lateral cusps. The crown sits horizontally on the massive neck, and reaches 0.4 mm in length and 0.25 mm in width. The pedicle base outline is curved, multipetaloid shaped (Fig 3O).

The longitudinal and transversally sectioned dermal denticles of euselachian [VU-ICH-KAR-014, VU-ICH-KAR-016] reveals the presence of a thin layer of enameloid covering the top surface of the crown (Fig 4A_2_–A_5_). This layer has a homogeneous thickness of about 0.005 - 0.0005 mm and gives the appearance of been composed loose bundles of enameloid oriented perpendicular to the crown surfaces. This could better appreciate in the longitudinal section (Fig 4B_2_–B_3_), whereas the poor preservation of the dermal denticles sectioned transversally only allow us to confirm the presence of the enameloid layer, but not the organization of it. This organization has also been described in euselachian dermal denticles of some Mesozoic Hybodontiformes and living neoselachian [33]; although in the dermal denticles of the neoselachian there appears to be two “sublayers”, with the outer sublayer composed of compacted crystals and the inner sublayer with crystals showing a preferred orientation perpendicular to the crown surface [33].

**Fig 4.**
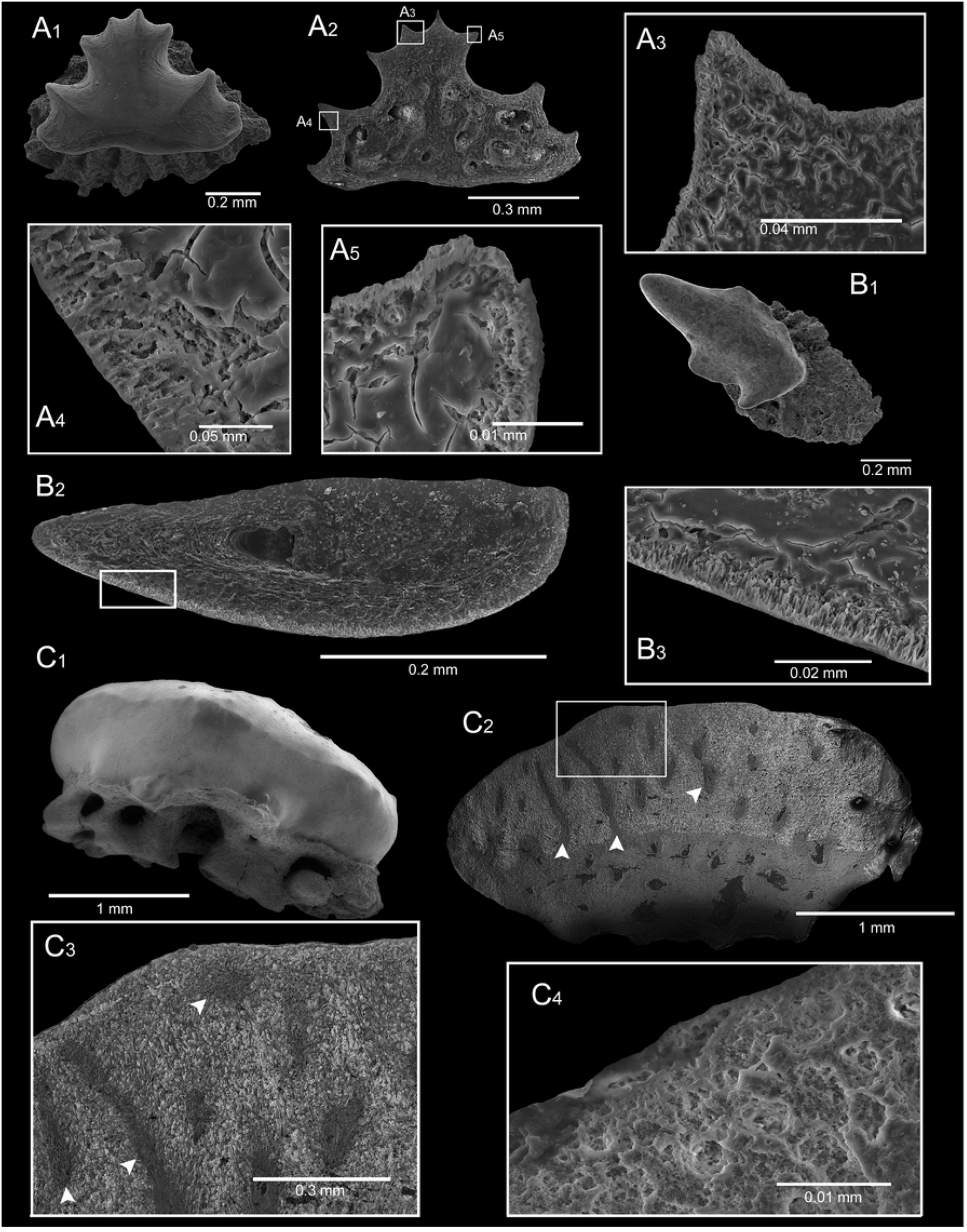
The histology of euselachian type dermal denticles and *Helodus* sp. tooth [VU-ICH-KAR-012] from Karpėnai quarry. (A) VU-ICH-KAR-014 from KAR-7 sample. (A_1_) general view of dermal denticle; (A_2_) general view of the longitudinally sectioned of denticle; (A_3_–A_5_) enameloid covering the top surface of the crown. (B) VU-ICH-KAR-016 from KAR-11 sample. (B_1_) general view of dermal denticle; (B_2_) general view of the longitudinally sectioned of denticle; (B_3_) enameloid covering the top surface of the crown. (C_1_) the entire isolated fossil; (C_2_) the longitudinal sectioned tooth with the well-differentiated cap of enameloid and distinct enameloid-dentine junction (EDJ); (C_3_) visualized in details the penetrating canals of enameloid layer marked by white arrows pointer; (C_4_) visualized in details a randomly arranged enameloid crystallites. White rectangles represent an outer layer of the crown in details.

The inner structure of euselachian type dermal denticle based on 3D-data is briefly described here [VU-ICH-KAR-023] (Fig 5A). The complete vascular canal system can be divided into two major types, based on their topological position in the studied fossil [50]. First type is placed on the neck and represented by five elongate, narrow and vertical canals which are oriented almost perpendicular to the denticle base - a conventional pattern (Fig 5A_5_–A_6_,A_8_–A_9_,A_11_–A_12_). It formed ‘bridge-like’ structure, creating a junction between more complicated developed vascular system of the crown (Fig 5A_2_–A_3_) and base with a complex geometry – a second type. In general, the canal system of euselachian dermal denticle has interconnected system without any distinctly isolated details.

**Fig 5.**
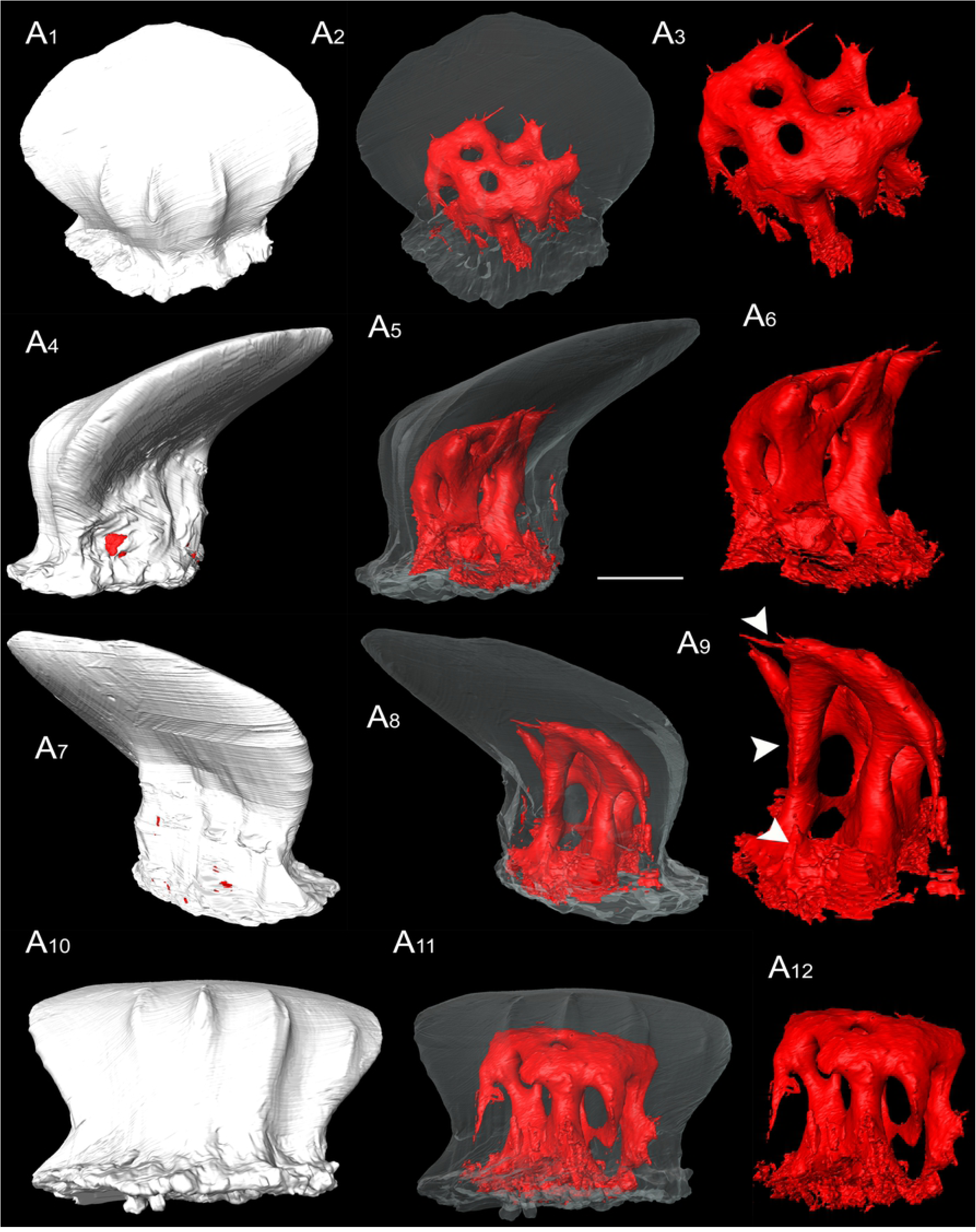
The 3D scanned euselachian type dermal denticle [VU-ICH-KAR-023] from KAR-4 sample, with details of the bone morphology and vascular system from Karpėnai quarry. (A_1_) external occlusal view; (A_2_, A_5_, A_8_, A_11_) shade bone; (A_3_, A_6_, A_9_, A_12_) vascular system with transparent bone contour; (A_4_) lingual-lateral view; (A_7_) oblique labial-lateral view; (A_10_) occlusal view. Major types of inner structure showed by white arrows. Scale bar equals 0.1 mm.

### Morphotype 6

Rare type of crown surface in studied quarry (Fig 3Q_1_–Q_2_). Mirror reflect ‘twins’ crown has many convex lateral short ridges and two long mesial ridges which reached the posterior margin (Fig 3Q_1_). The crown sits vertically on its base, creating ∼90° between these axes, thereby missing a neck considered to be an important morphological feature of this type. The subcrown is smooth, without any ornament (Fig 3Q_2_). The pedicle base outline is roundish, ellipse-shaped with strongly concave base, canal openings are not observed. The dermal denticle reaches 0.3 mm width and 0.2 mm length. Similar, wide opening for the pulp cavity scales are observed in Devonian thelodonts [51].

## Remarks

In general, this dermal denticles collection is clearly similar to the other euselachian morphotype dermal denticles which have already been described from the Naujoji Akmenė Formation (Lopingian) of Karpėnai quarry in the northern Lithuania [25].

Order HELODONTIFORMES Patterson, 1965 [52]

Family HELODONTIDAE Patterson, 1965 [52]

Genus *HELODUS* Agassiz, 1838 [41]

**Type Species**—*Helodus simplex* Agassiz, 1838 [41]

*HELODUS* sp.

(Figs 4C,6)

**Fig 6.**
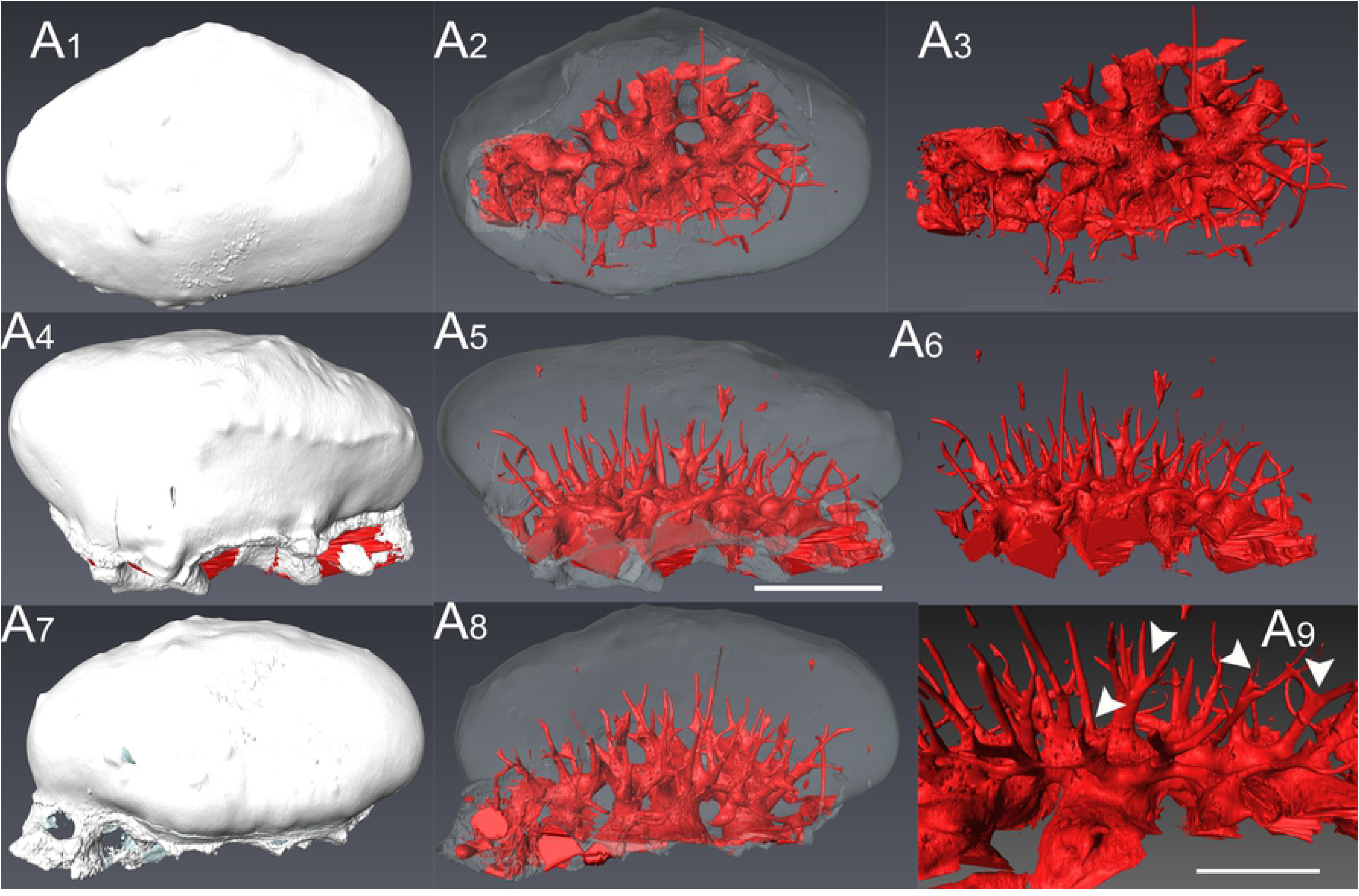
The 3D model of scanned *Helodus* sp. tooth [VU-ICH-KAR-012] from KAR-7 sample showing more details of the entire canal system from Karpėnai quarry. (A_1_) external view; (A_2_, A_5_, A_8_) shade bone; (A_3_, A_6_, A_9_) vascular system with transparent bone contour; (A_4_) lingual-lateral view; (A_7_) oblique labial-lateral view. Major types of pore cavities showed by white arrows. Scale bars equal: 1.0 mm (A_1_, A_2_, A_3_, A_4_, A_5_, A_6_, A_7_, A_8_) and 0.25 mm (A_9_).

## Material

Isolated well-preserved tooth from Karpėnai quarry, northern Lithuania [VU-ICH-KAR-012] and poorly-preserved tooth from Kūmas quarry, southern Latvia (not pictured here).

## Description

The poorly-preserved isolated tooth crown fragment is smooth, massive, has convex roundish outline with single sharper cusp in lingual margin. The tooth fragment reaches 0.7 mm in length and 0.5 mm in width.

The longitudinal section of the single *Helodus* sp. tooth reveals the presence of a well-differentiated cap of enameloid that coats the surface of it, with a quite distinct enameloid-dentine junction (EDJ) (Fig 4C_1_–C_2_). This monolayer consists of single crystallite enameloid (SCE) lacking any kind of organization, with the enameloid crystallites randomly arranged (Fig 4C_4_). The enameloid crystallites are round or subround (Fig 4C_4_), and the EDJ is clear. Penetrating the enameloid layer there are canals issuing from the dentine below (Fig 4C_2_) nearly reaching the surface of the tooth (Fig 4C_3_). This organization is similar to the one that was found in the tooth plate of a Carboniferous *Helodus* from England, although the enamloid layer in our tooth is thicker [53].

The 3D model of *Helodus* sp. tooth (Fig 6A_1_, A_4_, A_7_) showing a complete vascular system (Fig 6A_2_–A_3_,A_5_–A_6_,A_8_) which consists of some major types of pore cavities (white arrows Fig 6A_9_). Most of the cavities are single and isolated from other cavities. Another type is paired, with a small shared basal part close to the junction between the pore cavities and canal. A third type shares a larger basal portion and the two cavities diverge at a higher position. The last type shows an extreme condition with two pore cavities diverging at a position close to the surface of the tooth [50]. The vertical, narrow and elongate pore cavities are oriented oblique perpendicular to the tooth base.

## Remarks

The tooth has same morphological features as well-preserved Helodus-like tooth [VU-ICH-KAR-012] from Naujoji Akmenė Formation of late Permian in north Lithuania [25]. Also, the general shape of this tooth is comparable to a description of a *Helodus* tooth from Permian of western Australia [26]. In this paper, we studied it in more detail by investigating inner tissue composition using hand-made ground section of this tooth. Moreover, this fossil was scanned by the synchrotron and its slice images were imported to the Avizo 3.1 software for the 3D reconstruction model.

Superclass OSTEICHTHYES Huxley, 1880 [37]

The most common osteichthyan fossils in the Kūmas quarry are represented by their isolated scales and teeth (Fig 7-8). Rare in these assemblages are poorly preserved bony fish vertebral centra, (Fig 7N) and possible otoliths (not pictured here) and a single unidentified actinopterygian tooth plate (Fig 7O).

Class ACTINOPTERYGII Cope, 1887 [55]

Order PALAEONISCIFORMES Hay, 1902 [46]

Family PALAEONISCIDAE Vogt, 1852 [56]

PALAEONISCIDAE indet.

(Fig 7)

**Fig 7.**
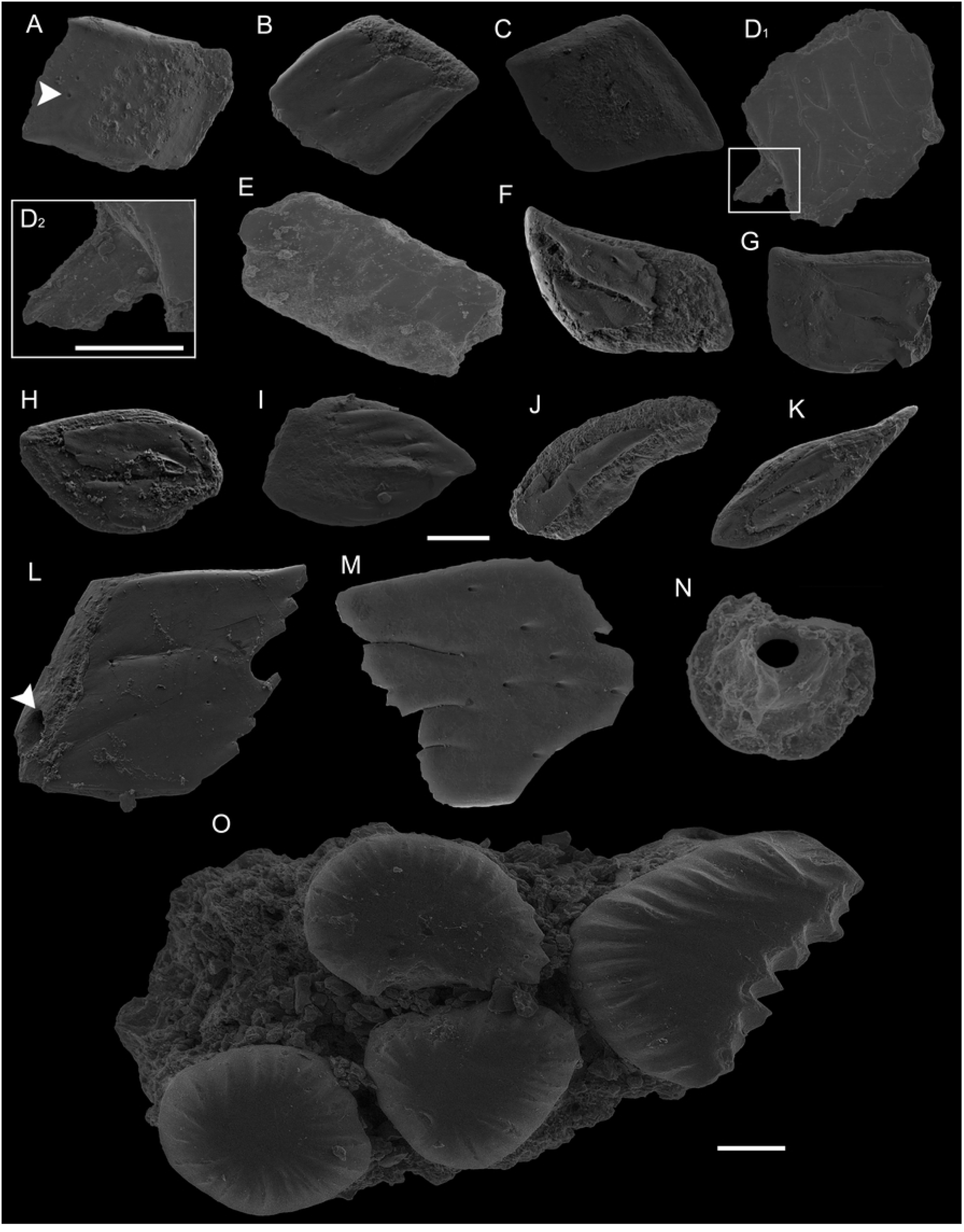
SEM images of isolated *Palaeoniscus* scales from KUM-2 sample, an actinopterygian unidentified vertebral centra from KUM-2 sample and actinopterygian teeth plate from KUM-3 sample of Kūmas quarry in general view. (A) VU-ICH-KUM-018; (B) VU-ICH-KUM-019; (C) VU-ICH-KUM-020; (D) VU-ICH-KUM-021; (D_1_) a general view; (D_2_) a detailed peg view; (E) VU-ICH-KUM-022; (F) VU-ICH-KUM-023; (G) VU-ICH-KUM-024; (H) VU-ICH-KUM-025; (I) VU-ICH-KUM-026; (J) VU-ICH-KUM-027; (K) VU-ICH-KUM-028; (L) VU-ICH-KUM-029; (M) VU-ICH-KUM-030; (N) VU-ICH-KUM-058; (O) VU-ICH-KUM-059. White arrow showed a canal opening on the top of crown (A) and a wide foramen (L). Scale bars equal: 0.1 mm (A, B, C, D_1_, E, F, G, H, I, J, K, L, M, N), 0.05 mm (D_2_) and 0.1 mm (O).

## Material

211 scales with well-preserved ganoine outer layer were found in the in Kūmas quarry, southern Latvia. All these scales were classified into 6 morphotypes based on their morphological features, which are shown in the 13 chosen samples [VU-ICH-KUM-018– VU-ICH-KUM-030].

## Description

The scales were identified to the specific fish body sectors as anterior-mid lateral flank, posterior lateral flank, ventral flank, pectoral peduncle, caudal fulcrum and unknown location scales based on its morphological features [57–59]. As a result, *Palaeoniscus* sp. scales can be divided into six morphotypes.

### Morphotype 1

These scales belong to posterior lateral frank and caudal peduncle fish body sectors (Fig 7A-D). It is rhombic or diamond shaped, sometimes is gently convex in central part (Fig 7A,C), but more common flat crown. Examination of these scales with SEM was disclosed numerous small, roundish-shape microtubercles in the outer ganoine layer. The crown is smooth, but some scales have fine, short grooves ornament which may have a direction from anterior to posterior side of the scale margins (Fig 7B,D). Only one of the specimens shows delicate perpendicular striation. The scales have some notable canal openings on the top of crown (white arrow Fig 7A). An angle location between margins indicates two obtuse for the dorsal, ventral and two acute for anterior, posterior corners. Rare case when all four angles of the scale are almost right (∼90°). The crown diagonal length reaches 0.6 – 1.5 mm. The base is thick and convex with a defined peg or without a peg. Such scales were found and identified as Haplolepidae family according to the similar rhomboidal scale from a haplolepid with a ganoine-covered outer surface and sinuous narrow grooves which were found in the Rader Limestone Member of the Bell Canyon Formation, Capitanian in West Texas, USA [60]. Moreover, similar *Gyrolepis* sp. rhomboidal scale was found in the North-Sudetic Basin, from Middle Triassic of Poland [60].

### Morphotype 2

The anterior-mid lateral flank scale (Fig 7E). It is an elongated rhombic shaped scale, with the right angle of all four corners and bad-preserved peg-and-socket. The crown ganoin is ornamented by several transverse inter-ridge grooves over the middle part of the surface. The base is slightly narrow and flat. The scale is 1.8 mm in length while a width reaches 0.7 mm. Notice: only single specimen of this morphotype was found.

### Morphotype 3

Scales of this type are located on the ventral flank sector [59] (Fig 7 F-G). It is generally elongated (Fig 7F); some scales have a square shaped (Fig 7G), thick, with slightly twisted posterior corner and other obtuse or almost right angles. The ganoine outer tissue has microtubercles pattern and covered up to about 0.75 mm of the scale depressed field. The crown has some canal openings and short, narrow concave ridges ornament which may has a direction from anterior to posterior side of the scale margins. The length reaches 1.0 mm and width is 0.4-0.6 mm.

### Morphotype 4

These scales belong to the pectoral peduncle sector (Fig 7H-I). They are small, elongated, almond shaped made by acute angle of posterior corner and slightly obtuse anterior corner. Their posterior and dorsal margins considerably longer than their anterior and ventral margins. The crown field constitutes of 3/4 ganoine tissue, with roundish microtubercles pattern (Fig 7H), some scales have overlapped wide convex ridges (Fig 7I), grew up in the anterior-posterior direction. There is no socket-and-peg on the thick base. Such scales reach 0.6-0.8 mm in length and 0.4-0.5 mm in width.

### Morphotype 5

This type scales located on the caudal fulcrum sector (Fig 7J-K). They are generally small, thick, rod-like, or more precisely, extremely elongated shape with flat, slightly massive crown, 3/4 covered by ganoine tissue. The crown surface sculpted by tiny, roundish microtubercles, has no ornament. Such scales antero-dorsal corner is tapering, its margins looks convex while the postero-ventral corner is acute, lightly concave, much narrow, ganoine layer is absent. The scale length reaches 1.0 mm while width only 0.2-0.3 mm.

### Morphotype 6

This type of scales belonged to undetermined fish body sector, because material is not completely preserved (Fig 7L-M). Such samples composed out of the straight dorsal, ventral and anterior margins while entire posterior margin is pectinated, has several narrow serrations. The crown field is covered by ganoine tissue with tiny, roundish microtubercles pattern. Some scales are smooth, showing flask-shaped pore cavities visible through translucent ganoine layer (Fig 7M). Sometimes, the anterior margin is thick, slightly massive with significant, wide foramen of vascular system (white arrow Fig 7L). The scales length reaches 1.3-1.5 mm and 1.0 mm in width. Similar shaped and described scales were obtained in Elonichthydae family based on its features of the microstructure on the outer surface from the Black Shale Horizon of the Syřenov Formation in Czech Republic [62].

## Remarks

Most of the scales belong to the Palaeonisciformes order based on a comparison with Palaeoniscoid scales [63], Palaeonisciformes-related material [62] and its description [64]. Less variable, similar actinopterygian scales assemblage was found in Karpėnai quarry, northern Lithuania [25].

Family PALAEONISCIDAE Vogt, 1852 [56]

PALAEONISCIDAE indet.

(Figs 8 A-M)

**Fig 8.**
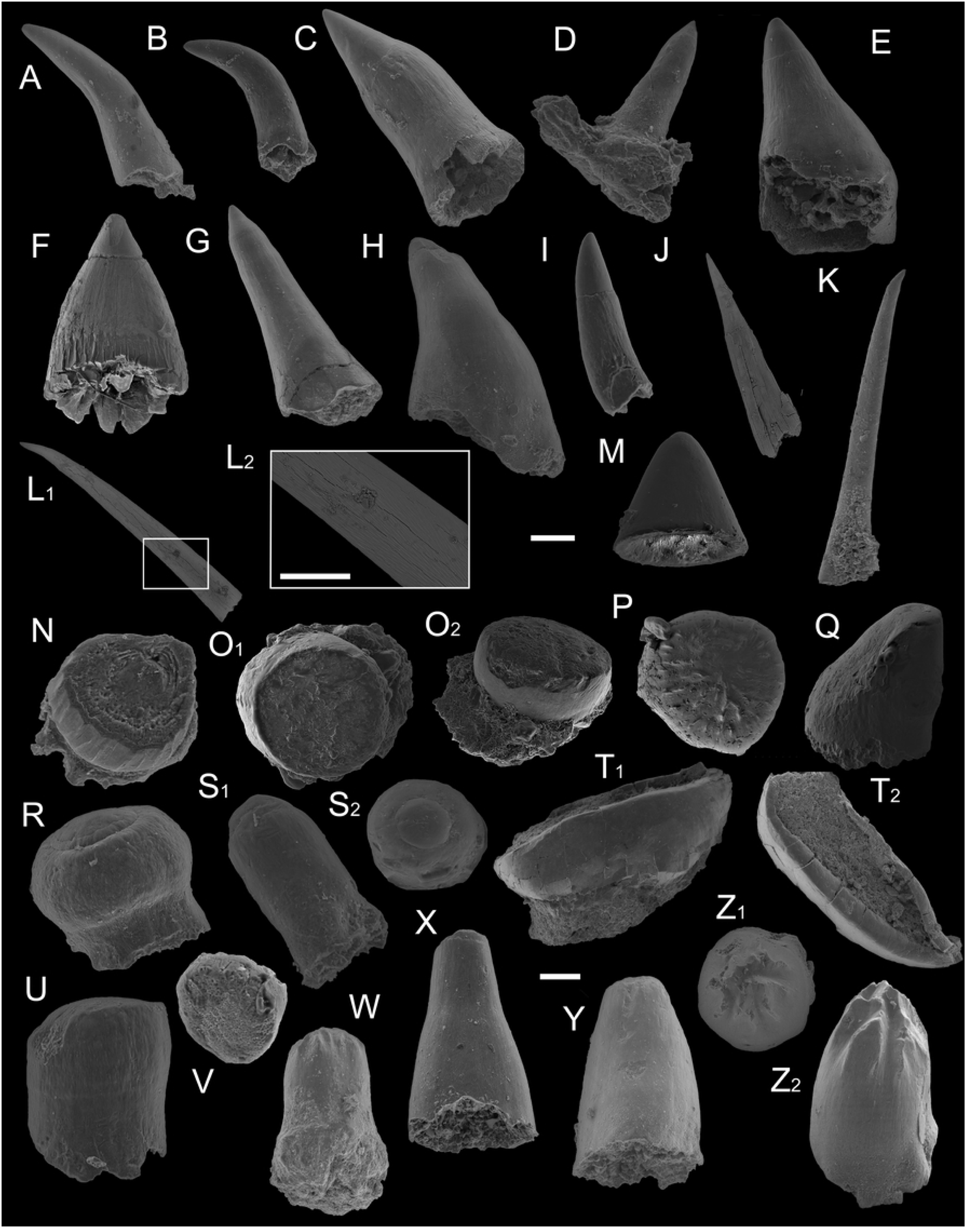
SEM images of the different shaped actinopterygian teeth actinopterygian teeth assemblage from Kūmas quarry. (A) VU-ICH-KUM-031 from KUM-4 sample. (B) VU-ICH-KUM-032 from KUM-3 sample. (C) VU-ICH-KUM-033 from KUM-3 sample. (D) VU-ICH-KUM-034, with preserved fragment of the base from KUM-3 sample. (E) VU-ICH-KUM-035 from KUM-4 sample. (F) VU-ICH-KUM-036 from KUM-1 sample. (G) VU-ICH-KUM-037 from KUM-4 sample. (H) VU-ICH-KUM-038 from KUM-3 sample. (I) VU-ICH-KUM-039 from KUM-10 sample. (J) VU-ICH-KUM-040 from KUM-2 sample. (K) VU-ICH-KUM-041 from KUM-3 sample. (L) VU-ICH-KUM-042 from KUM-4 sample; (L_1_) general view; (L_2_) microtubercles structure of tooth. (M) VU-ICH-KUM-043 from KUM-9 sample, enameloid cap; lateral view. ?*Platysomus* sp. teeth. (N) VU-ICH-KUM-044 in oblique occlusal view from KUM-9 sample. (O) VU-ICH-KUM-045 from KUM-3 sample; (O_1_) in occlusal view; (O_2_) in lateral view. (P-Z) indeterminate actinopterygian teeth. (P) VU-ICH-KUM-046 in occlusal view from KUM-5 sample. (Q) VU-ICH-KUM-047 in lateral view from KUM-3 sample. (R) VU-ICH-KUM-048 in lateral view from KUM-3 sample. (S) VU-ICH-KUM-049 from KUM-4 sample, (S_1_) in lateral view; (S_2_) in occlusal view. (T) VU-ICH-KUM-050 from KUM-5 sample, (T_1_) in lateral view; (T_2_) in occlusal view. (U) VU-ICH-KUM-051 in lateral view from KUM-4 sample. (V) VU-ICH-KUM-052 in oblique occlusal view from KUM-4 sample. (W) VU-ICH-KUM-053 in lateral view from KUM-4 sample. (X) VU-ICH-KUM-054 in lateral view from KUM-4 sample. (Y) VU-ICH-KUM-055 in oblique lateral view from KUM-4 sample. (Z) VU-ICH-KUM-056 from KUM-3 sample; (Z_1_) in oblique lateral view; (Z_2_) in occlusal view. Scale bars equal: 0.1 mm (A-L_1_, M-S,U-Z), 0.05 mm (L_2_) and 0.3 m (T).

## Material

1671 well preserved isolated teeth in Kūmas quarry, southern Latvia. Teeth are represented by the 13 different pictured microremains [VU-ICH-KUM-031–VU-ICH-KUM-043].

## Description

Varied assemblage of the conical, thin, light amber coloured teeth. Most of the teeth are slender, smooth, without distinct visible sculpture, although microtubercles are well developed (Fig 8D,G,I-M). They covered the whole surface of the tooth, with the exception of the acrodin cap. Microtubercles are proximo-distally elongated, narrow and blend together with oblique rows (Fig 8L_2_). Such teeth reach from 0.3 mm till 4.0 mm in length while the width is 0.09 – 0.3 mm. Common teeth are straight with the wider dentin cones and its base (Fig 8C,E,F,H). These teeth have not any distinct sculptures and could reach a maximum width, length measurements of 0.6 mm. Very rare teeth are elongate, smooth, curved ‘horn-like’ shape with slightly blunt transparent acrodin cap (Fig 8A-B). Such teeth reach 0.3-0.6 mm in length and 0.05-0.2 mm in width.

The SEM analysis of the teeth (Fig 9A_1_, B_1_) reveals an enameloid layer surrendering a dentine core with a central pulp cavity (Fig 9A_2_). The enameloid extends only over the apical part of the tooth, attaining its maximum thickness at the base of the cap where it reaches 70 µm (Fig 9A_3_). Lingually and labially, the enameloid layer is reduced to 5 µm (Fig 9A_4_–A_5_). The hypermineralised layer is composed of single crystallites with different organization [64], depending on the part of the layer in which they are embedded. Lingually and labially, the layer is composed of individual crystallites parallel to each other and oriented perpendicular to the tooth surface, without any kind of higher form of organization (Fig 9A_5_). In the base of the cap, where the layer is thicker, the enameloid crystallites are organised in loose bundles (Fig 9A_3_) that display a tendency to be oriented parallel to the tooth surface (Fig 9B_2_). The teeth do not show any kind of superficial layer, which may have been removed in the process of tooth wear. The enameloid-dentine junction (EDJ) is clearly distinguishable and regular.

**Fig 9.**
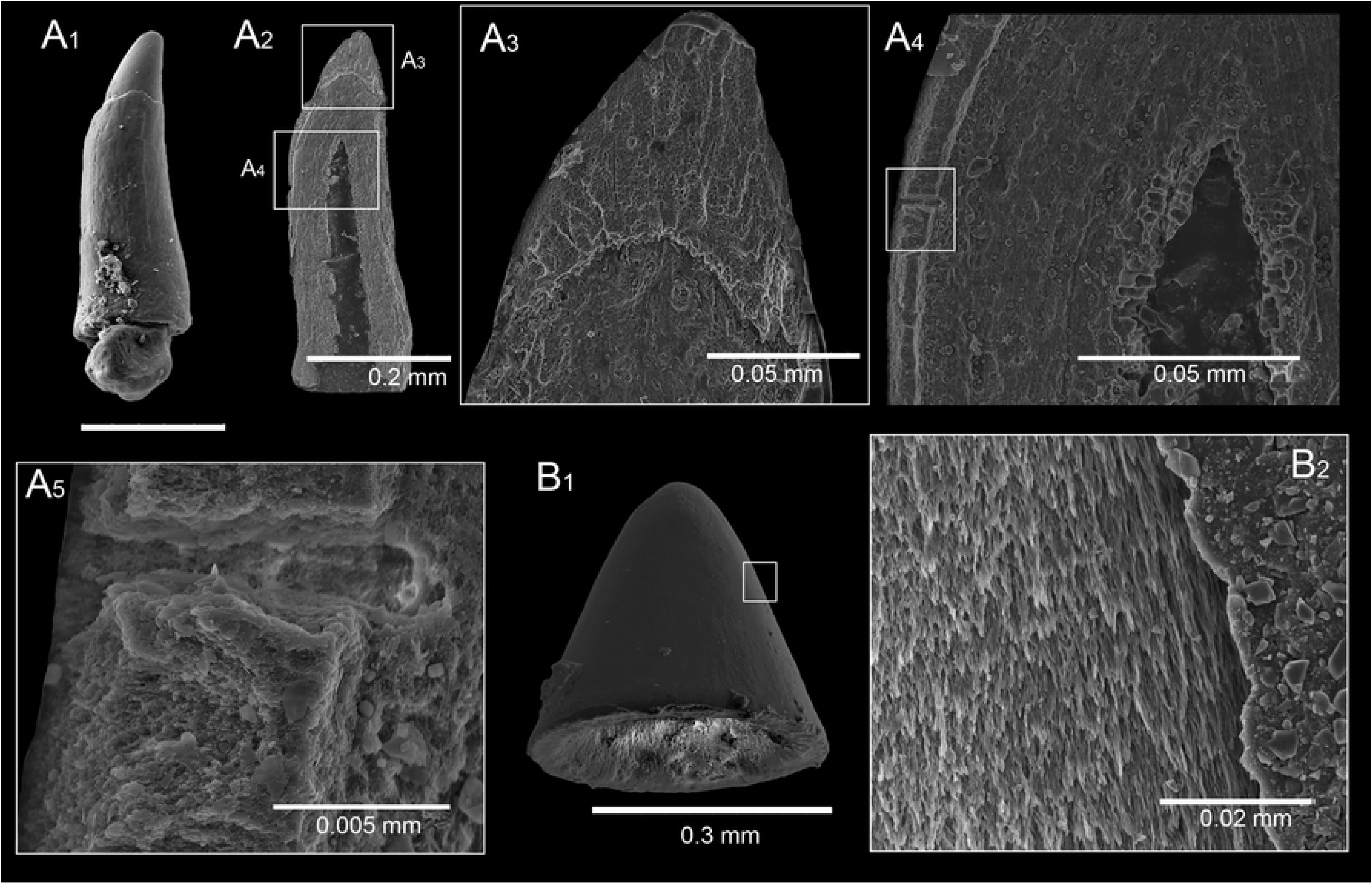
The histology of Palaeoniscus teeth from Kūmas quarry. (A_1_) general view of the tooth [VU-ICH-KUM-057] from KUM-2 sample; (A_2_) the entire longitudinal sectioned tooth showing enameloid layer, dentine core and central pulp cavity; (A_3_) an enameloid layer in details; (A_4_) a collar enamel (by Sasagawa et al., 2009); (A_5_) the individual crystallites parallel to each other and oriented perpendicular to the tooth surface. (B_1_) general view of the acrodine cap [VU-ICH-KUM-043] from KUM-9 sample; (B_2_) enameloid cap showing any crystallites orientation.

## Remarks

Similar smooth, straight, long teeth with microtubercles were identified as *Elonichthys* sp. from Krkonoše Piedmont Basin of the Stephanian in Czech Republic [62]. Also, the same assemblage with less variety was obtained from Naujoji Akmenė Formation, late Permian in Lithuania [25].

Family PLATYSOMIDAE Young, 1866 [66]

Genus *PLATYSOMUS* Agassiz, 1843 [42]

**Type Species**—*Platysomus gibbosus* Blainville, 1818 [67]

*?PLATYSOMUS* sp.

(Fig 8N-O)

## Material

Five isolated teeth were found in Kūmas quarry, southern Latvia. Only two teeth pictured and represented here, show morphological differences [VU-ICH-KUM-044; VU-ICH-KUM-045].

## Description

The teeth have a rod-shaped, short cylinder outline with flat apex (note: tooth cap was probably lost?) (Fig 8N-O_1_). Some teeth have many vertical, convex ridges which covered an entire lateral margin and with visible enameloid, dentine tissues inner structure in the cross section in apical view (Fig 8N). While other teeth have only well-developed vertically elongate, narrow microtubercles in lateral view with smooth rubbed apex (Fig 8O_2_). The tooth reaches a maximum of 0.5 mm in diameter and 0.2 mm in height.

## Remarks

These teeth could be assigned to *Platysomus* or *Kargalichthys*-like tooth based on the description of palaeoniscoid assemblage from the Upper Permian of the European part of Russia [68]. Also, it similar to *Lepidotes*-like tooth based on the SEM photograph from Late Jurassic of Uruguay [69]. However, the earliest Semionotiformes are only known from the Triassic [70].

### Actinopterygian indeterminate microremains

The actinopterygian indeterminate microremains are represented here by isolated teeth, and dental remain and divided into 4 groups (A, B, C, D) based on its different morphological characteristics.

Group A, consists of eight isolated teeth [VU-ICH-KUM-046–VU-ICH-KUM-047]. The seven teeth are semispherical, convex, with circular outline shaped in apical view. The entire crown has features of the irregular and radiating ridges which do not join in the central part of the cap (Fig 8P). Microtubercles are well developed and consist out of proximo-distally elongated, narrow and blend together with oblique rows. Such teeth reach a maximum diameter of 1.2 mm and 0.5 mm in height. Single Pycnodontidae tooth has an asymmetrical conical crown with straight, slightly convex labial-lateral margin and a relatively narrow occlusional platform confluent with the strongly concave lingual face (Fig 8Q). This tooth reaches 1.0 mm in height and 0.7 mm in base width. The low profile and circular outline of this tooth suggest pycnodont affinities. A similar circular tooth with radiating margin ridges has been illustrated and identified as cf. *Anomoeodus* from the Sao Khua Formation of the Cretaceous at Phu Phan Thong in Thailand [71]. Moreover, the studied tooth was found the morphological similarities to Pycnodontiformes indet. from the Lower Cretaceous in Tunisia [72]. Identic-shaped teeth were earlier found from Naujoji Akmenė Formation of Upper Permian in Lithuania [25]. Such robust conical crowns tooth was found from the Upper Cretaceous in Bulgaria [73].

Group B, includes six crushing-type teeth and an isolated dental remain [VU-ICH-KUM-048–VU-ICH-KUM-050]. The crushing teeth represented here as globular (Fig 8R) and circular-rod (Fig 8S) shaped in lateral view, with convex, tiny, sometimes wide, hemispherical acrodine cap in apical view. The crown is smooth, without any distinct visible ornament. Only the roots are missing. Such teeth could reach 0.1-0.5 mm in diameter and 0.4-0.6 mm in height. While massive dental remain has a long axis parallel to the labial margin, narrow and bilaterally symmetrical shape with slightly depressed oval, thick, ring-like rim of the enameloid layer in apical view (Fig 8T_1_). The tooth crown is smooth, its central indent is shallow and filled by outer sediments (Fig 8T_2_). The base is broad and partly preserved. The length has reached 2.5 mm, width – 0.6 mm and height – 1.1 mm. Similar vomerine tooth plate of *Macropycnodon streckeri* was found in Juana Lopez Member of the Upper Cretaceous from New Mexico [74] and cf. *Coelodus* sp. *vomers* from Csehbánya Formation of the Upper Cretaceous in Hungary [75]. Moreover, crushed-type teeth were identified as aff. *Gyrodus?* sp. from Lower Creatceous of the Baltic Sea [76]. However, another globular and conical tooth with a little central ‘wart’ was assigned to *Lepidotes* sp. from the Middle Jurassic of the Grands Causses in southern France [77].

Group C has three isolated teeth [VU-ICH-KUM-051–VU-ICH-KUM-52]. Crushing dentition has spherical or styliform shaped in lateral view (Fig 8U) with circular outline, flat, smooth crown tip in apical view and without any visible ornament (Fig 8V). Such teeth reach 0.4-0.5 mm in height and 0.25 mm in diameter. These teeth are similar to the Lepisosteiformes order based on the fish findings, their images and descriptions from Wessex Formation of Lower Cretaceous in the Isle of Wight, southern England [78].

Group D includes several teeth [VU-ICH-KUM-053–VU-ICH-KUM-056]. Such a collection of the teeth is conical shaped and covered by microtubercles, which are proximo-distally elongated, narrow and blend together with oblique rows. The crown has features of the irregular and radiating ridges which creating blunt (Fig 8W) or slightly acute (Fig 8Z) central part of the cap. Sometimes the teeth cap devoid of any ornamentation (Fig 8X-Y). Such teeth reach 0.5 mm in height and 0.3 mm in diameter.

## Discussion

The results presented in this paper reveal a late Permian fish assemblage from Kūmas quarry locality of southern Latvia. The late Permian osteichthyans and chondrichthyans were dominant fish classes among marine and freshwater vertebrates all over the world [2–3]. The osteichthyan microremains in the studied sites consisted of the numerous isolated teeth and scales of Palaeoniscidae, Platysomidae, Haplolepidae and Elonichthydae, while chondrichthyans are represented by various dermal denticles and teeth of Euselachii, Helodontidae.

The Hybodontoidei remains and Euselachii dermal denticles are already known from Early Permian of Texas [79]; from Middle Permian of Arizona [80]; from Lopingian of Hydra Island, Greece [20]; from Kazanian Stage, Middle Permian of Russia [81]; and Naujoji Akmenė Formation, late Permian of Lithuania [25]. Also, the Hybodontidae dermal denticles were found under Permian-Triassic boundary in Zhejiang and Jiangxi provinces in South China [12–13]. *Helodus* sp. fragments have been found in Permian deposits of West Australia [26] and Naujoji Akmenė Formation of Lithuania [25]. Teeth of *Omanoselache* sp. are know from Khuff Formation of Sultanate of Oman [45]. *Acrodus* teeth crown were found in late Permian deposits of the central Iran [18].

The osteichthyan fishes had a very wide paleogeographical distribution during the Permian [3]. Diverse chondrichthyan and actinopterygian microremains are known from central Iranian locality of Baghuk Mountain, late Permian [18]. Whole Platysomidae skeletons and some isolated fragments have been found in the Upper Kazanian Stage, the Middle Permian, and Upper Permian of the East European Platform of Russia [16,83]; almost complete *Dorypterus* sp. was found in the Permian of England [84]; isolated shark and actinopterygian teeth, scales are known from El Jarillal Formation of Permian in Argentina [85]; *Blourugia* sp. actinopterygians were found in the late Permian deposits of the Beaufort Group, South Africa [8]; well-preserved ?*Platysomus*sp. and *Palaeoniscum* sp. skeletons have been found in the Lopingian Stage, the Upper Permian in northwest Germany [21]. *Elonichthys* sp. skeletons and skull fragments came from Permo-Carboniferous lakes strata in the Czech Republic and in the Saar-Nahe Basin in southwest Germany [86–89]; some *Elonichthys* sp. microremains have been found in Permian deposits in Norway [90]. *Polyacrodus* sp. teeth, symmoriiform denticles, euselachian scales, *Stethacanthulus meccaensis* tooth, Haplolepid scales, Elonichthyid scales, *Variolepis* sp. scales are known from the Middle Permian of West Texas, USA [60].

A similar late Permian assemblage of isolated chondrichthyan and osteichthyan fossils was earlier found and described in the Karpėnai quarry, northern Lithuania [25]. The comparison of fish fossil abundances from the roughly similar total samples (weighting ∼210 kg) of limestones between Karpėnai and Kūmas quarries reveal their significant difference. Hence, actinopterygian teeth are ∼120 times and actinopterygian scales are 23 times more abundant in Latvian quarry. The comparison of chondrichthyans microremains showed that dermal denticles are two times and their teeth are 17 times more abundant in the Kūmas quarry than the Karpėnai quarry of the same age that represents more distal environments. The difference in the influx of fresh water from the surrounding terrain towards the hypersaline Zechstein Basin possibly was a major factor that affected differences in the abundance of the fossil material between Kūmas locality which was much closer to the shoreline (and thus riverine source of water) and the Karpėnai locality, which was further offshore [91–92]. The Zechstein Basin was significantly restricted sea with sedimentation that resembled giant playa like sequences, where hypersaline waters in the central part gave rise to evaporites together with carbonates and more fresh water environments near shoreline were suitable to the development (precipitation) of limestones [29,93]. Therefore, current observation of the abundance and diversity gradient is congruent with the explanation that the major factor that determined higher richness of fossil ichthyofauna in the Kūmas locality compared to the southern Karpėnai locality is the salinity gradient. The distribution of anhydrites and gypsum that are abundant in the correlates of the Naujoji Akmenė Formation in central Lithuania and Kaliningrad Oblast of Russia [29], suggest that the salinity gradient increased in the south-western direction from the studied sites, thus probably physiologically restricting possible distribution of fish faunas. In addition to the salinity stress, ichthyofaunas which existed in the offshore parts of the Zechstein sea, should have experienced stronger cytotoxic effects of halogenated hydrocarbons. Halophilic bacteria - the major source of volatile halogenated hydrocarbons, thrive in modern hypersaline basins such as Kara Bogaz Gol, which are thought to be close environmental analogous to the hypersaline parts of the Zechstein sea [94]. Also, the variation in abundance of fish microremains in a stratigraphic succession could be explained by regional variations in fresh water input during episodic exposure events associated with regional or global sea-level fluctuations [92].

## Conclusion

The new chondrichthyan and osteichthyan material from late Permian of Kūmas quarry (southern Latvia) presented here adds a new data on the scarce ichthyofaunal fossil record of the northeastern Zechstein Basin margin.

The influx of fresh water could have affected the fossil abundance differences between more offshore Karpėnai locality and nearshore Kūmas quarry locality of Zechstein Basin. Well-bedded limestone of Naujoji Akmenė Formation is easily correlatable with Sātiņi, Kūmas, and Alši Formations limestones.

The numerous isolated fish microremains were found and identified as belonging to Palaeoniscidae, Platysomidae, Haplolepidae and Elonichthydae from the osteichthyan clade, while chondrichthyans are represented by various dermal denticles and teeth of the Euselachii, Helodontidae, ?*Acrodus* sp., and ?*Omanoselache* sp.

SEM imaging analysis revealed that late Permian primitive fish (chondrichthyans, actinopterygians) teeth possessed an outer homogeneous layer of SCE described by matrix of the randomly orientated (fluor-) hydroxyapatite crystallites with non-preferred orientation.

3D-models of the vascular system and bone of euselachian type dermal denticle and *Helodus* sp. tooth from late Permian were visualised here. The synchrotron segmentation technique demonstrated and provided important information about the analysed fossil microanatomical structure and virtual histology without damaging the fish microremains. Euselachian inner structure of the dermal denticle represented by five single vertical canals, which joined a more complicated developed vascular system of the crown and base with a complex geometry. *Helodus* sp. vascular system of tooth composed of four main cavities types: single and isolated from other cavities; paired, with a small shared basal part close to the junction between the pore cavities and canal; a larger basal portion and the two cavities diverge at a higher position; two pore cavities diverging at a position close to the surface of the tooth. This 3D-data will thus be essential for fully understanding of the fish fossils architectural internal canal diversity and external morphological features for comparative studies in the future.

## Acknowledgements

The authors thank J.-P. M. Hodnett and J. Fischer for reviewing the manuscript, the I. Celiņš (CEMEX company, Latvia) who accompanied us in Kūmas quarry and D. Šiupšinskas (Vilnius University, Lithuania) who helped to collect and safely transport samples. Also, we would like to thank H. Botella and C. Martínez-Pérez (University of Valencia, Spain) for scanning Lithuanian fossils using synchrotron in Switzerland and sharing their experiences in the tomography and how to create 3D models using Avizo 8.1 software.

## References

1. Wignall PB. The Worst of Times: How Life on Earth Survived Eighty Million Years of Extinctions. Princeton University Press. 2015.

2. Koot MB. Effects of the Late Permian mass extinction on chondrichthyan palaeobiodiversity and distribution patterns. Ph.D. dissertation, University of Plymouth, Plymouth, England. 2013, 859 p.

3. Romano C, Koot MB, Kogan I, Brayard A, Minikh AV, Brinkmann W, et al. Permian–Triassic Osteichthyes (bony fishes): diversity dynamics and body size evolution. Biological Reviews. 2016 Feb;91(1):106–47.

4. Sallan LC, Coates MI. End-Devonian extinction and a bottleneck in the early evolution of modern jawed vertebrates. Proceedings of the National Academy of Sciences. 2010;107(22):10131–5.

5. Campbell KS, Phuoc LD. A Late Permian actinopterygian fish from Australia. Palaeontology. 1983;26(Part 1):33–70.

6. Bender PA. First documentation of similar Late Permian actinopterygian fish from Australia and South Africa. Records of the Western Australian Museum Supplement. 1999;57:183–9.

7. Bender PA. A new actinopterygian fish species from the Late Permian Beaufort Group, South Africa. Palaeontologia Africana. 2001;37:25–40.

8. Bender PA. A new deep-bodied Late Permian actinopterygian fish from the Beau fort Group, South Africa. Palaeontologia Africana. 2005;41:7–22.

9. De Figueiredo FJ, Carvalho BC. A new Actinopterygian fish from the Late Permian of the Paraná Basin, southern Brazil. Arquivos do Museu Nacional. 2004;62(4):531–47.

10. Richter M. First record of Eugeneodontiformes (Chondrichthyes: Elasmobranchii) from the Paraná Basin, Late Permian of Brazil. Paleontologia: cenários de vida. Interciéncía Ltda., Rio de Janeiro. 2007;1:149–56.

11. Poplin C, Wang NC, Richter M, Smith M. An enigmatic actinopterygian (Pisces: Osteichthyes) from the Upper Permian of China. Zoological Journal of the Linnean Society. 1991;103(1):1–20.

12. Wang N, Zhu X, Jin F, Wang W. Chondrichthyan microremains under Permian-Triassic boundary both in Zhejiang and Jiangxi provinces, China-Fifth report on the fish sequence study near the Permian-Triassic boundary in South China. Vertebrata Palasiatica. 2007;45(1):29. Chinese.

13. Wang NZ, Zhang X, Zhu M, Zhao WJ. A new articulated hybodontoid from Late Permian of northwestern China. Acta Zoologica. 2009;90:159–70.

14. Goto M. Palaeozoic and early Mesozoic fish faunas of the Japanese Islands. Island Arc. 1994 Dec;3(4):247–54.

15. Minikh AV. A new paleoniscoid fish from Upper Permian of East-European Platform. Paleontologicheskiy Zhurnal. 1990;3:71–76. Russian.

16. Esin DN. New species of deep-bodied actinopterygians (Platysomidae) from the Upper Permian of the East European Platform. PALEONTOLOGICHESKII ZHURNAL. 1993;(3):128–32. Russian].

17. Esin D. Ontogenetic development of the squamation in some Palaeoniscoid fishes. Bulletin du Muséum national d’Histoire naturelle, 4ème série–section C–Sciences de la Terre, Paléontologie, Géologie, Minéralogie. 1995;17(1-4):227–34. Russian.

18. Hampe O, Hairapetian V, Dorka M, Witzmann FL, Akbari AM, Korn D. A first late Permian fish fauna from Baghuk Mountain (Neo-Tethyan shelf, central Iran). Bulletin of Geosciences. 2013 Jan 1;88(1):1–20.

19. Nakazawa K, Kapoor HM, Ishii KI, Bando Y, Okimura Y, Tokuoka T. The upper Permian and the lower Triassic in Kashmir, India. Memoirs of the Faculty of Sciences, Kyoto University, Geology-Minerology. 1975;42:1–106.

20. Argyriou T, Romano C, Carrillo-Briceño JD, Brosse M, Hofmann R. The oldest record of gnathostome fossils from Greece: Chondrichthyes from the Lopingian of Hydra Island. Palaeontologia Electronica. 2017;20(1):1–9.

21. Diedrich CG. A coelacanthid-rich site at Hasbergen (NW Germany): taphonomy and palaeoenvironment of a first systematic excavation in the Kupferschiefer (Upper Permian, Lopingian). Palaeobiodiversity and Palaeoenvironments. 2009;89(1-2):67–94.

22. King W. The Permian fossils of England. Monograph of the Palaeontographical Society, London, England; 1850; 1–253 p.

23. Nielsen E. On new or little known Edestidae from the Permian and Triassic of East Greenland. De danske ekspeditioner til Ostgronland, København. 1952, 55 p.

24. Kazmierczak JO. Morphology and palaeoecology of the Productid horridonia horrida (Sowerby) from zechstein of Poland. Acta Palaeontologica Polonica. 1967;12(2).

25. Dankina D, Chahud A, Radzevičius S, Spiridonov A. The first microfossil record of ichthyofauna from the Naujoji Akmenė Formation (Lopingian), Karpėnai Quarry, North Lithuania. Geological Quarterly. 2017;61(3):602–10.

26. Teichert C. Bradyodont sharks in the Permian of Western Australia. American Journal of Science. 1943 Sep 1;241(9):543–52.

27. Zeeh S, Becker F, Heggemann H. Dedolomitization by meteoric fluids: the Korbach fissure of the Hessian Zechstein basin, Germany. Journal of Geochemical Exploration. 2000;69:173–6.

28. Raczyński P, Biernacka J. Zechstein in Lithuanian–Latvian Border Region. Geologija. 2014 Aug 20;56(2).

29. Paškevičius J. The Geology of the Baltic Republics. Vilnius University and Geological Survey of Lithuania, Vilnius, Lithuania. 1997, 387 p.

30. Van Wees JD, Stephenson RA, Ziegler PA, Bayer U, McCann T, Dadlez R, Gaupp R, Narkiewicz M, Bitzer F, Scheck M. On the origin of the southern Permian Basin, Central Europe. Marine and Petroleum Geology. 2000 Jan 1;17(1):43–59.

31. Kuršs V, Savvaitova L. Permian limestones of Latvia. Zinatne, Riga, Latvia, 1986; 94 p. Russian.

32. Gailīte LI, Kuršs V, Lukševiča L, Lukševičs E, Pomeranceva R, Savaitova L, et al. Legends for geological maps of Latvian bedrock. State Geological Survey, Riga; 2000. 101 p.

33. Jeppsson L, Anehus R, Fredholm D. The optimal acetate buffered acetic acid technique for extracting phosphatic fossils. Journal of Paleontology. 1999 Sep;73(5):964–72.

34. Manzanares E, Plá C, Martínez-Pérez C, Rasskin D, Botella H. The enameloid microstructure of euselachian (Chondrichthyes) scales. Paleontological Journal. 2014;48(10):1060–6.

35. Abel RL, Laurini CR, Richter M. A palaeobiologist’s guide to’virtual’micro-CT preparation. Palaeontologia Electronica. 2012;15(2).

36. Ivanov AO, Nilov SP. Microtomographic research of the vascularization system in the teeth of Palaeozoic sharks. In Bruker microCT User Meeting 2016.

37. Böttinger M, Meier-Fleischer K, Ulmen C. Tutorial: Interactive 3D Visualization in Earth System Research with Avizo Green 8.0. DKRZ/KlimaCampus Hamburg. 2013.135 p.

38. Huxley TH. On the application of the laws of evolution to the arrangement of the Vertebrata and more particularly of the Mammalia. InProceedings of the Zoological Society of London 1880 (Vol. 1880, pp. 649–662).

39. Bonaparte CL. Iconographia della fauna Italica per le quattro classi degli animali vertebrati, 3rd ed. Rome: 1832–1831.

40. Patterson C. British Wealden Sharks. Bulletin of the British Museum (Natural History). 1966;11:283–350.

41. Casier E. Contributions à l’étude des poissons fossiles de la Belgique. XII - Sélaciens et Holocéphales sinémuriens de la Province de Luxembourg. Bulletin de l’Institut Royal des Sciences Naturelles de Belgique. 1959;38(8):1–35

42. Agassiz L. Recherches sur les poisons fossils. 5th ed. Imprimerie de Petitpierre, Neuchâtel; 1833-1844, 1420 p.

43. Cuny G, Liard R, Deesri U, Liard T, Khamha S, Suteethorn V. Shark faunas from the Late Jurassic—Early Cretaceous of northeastern Thailand. Paläontologische Zeitschrift. 2014;88(3):309–28.

44. Błażejowski B. Shark teeth from the Lower Triassic of Spitsbergen and their histology. Polish Polar Research. 2004;25(2):153–67.

45. Koot MB, Cuny G, Tintori A, Twitchett RJ. A new diverse shark fauna from the Wordian (Middle Permian) Khuff Formation in the interior Haushi-Huqf area, Sultanate of Oman. Palaeontology. 2013 Mar;56(2):303–43.

46. Hay OP. Bibliography and catalogue of the fossil vertebrata of North America. Bulletin of the United States Geological Survey, 1901; p. 1–868.

47. Arratia G, Kriwet J, Heinrich WD. Selachians and actinopterygians from the Upper Jurassic of Tendaguru, Tanzania. Fossil Record. 2002;5(1):207–30.

48. Johns MJ. Diagnostic pedicle features of Middle and Late Triassic elasmobranch scales from northeastern British Columbia, Canada. Micropaleontology. 1996 Dec 1:335–50.

49. Ivanov AO, Nestell GP, Nestell MK. Fish assemblage from the Capitanian (Middle Permian) of Theapache Mountains, West Texas, USA. The Carboniferous-Permian Transition: Bulletin 60. 2013;60:152.

50. Reif WE. Types of morphogenesis of the dermal skeleton in fossil sharks. Paläontologische Zeitschrift. 1978;52:110–128.

51. Qu Q, Blom H, Sanchez S, Ahlberg P. Three-dimensional virtual histology of silurian osteostracan scales revealed by synchrotron radiation microtomography. Journal of morphology. 2015;276(8):873–88.

52. Örvig T. Thelodont scales from the Grey Hoek Formation of Andrée Land, Spitsbergen. Norsk Geologisk Tidsskrift. 1969;49(4):387–401.

53. Patterson C. The phylogeny of the chimaeroids. Philosophical Transactions of the Royal Society of London. 1965. B249:101–219.

54. Gillis JA, Donoghue PC. The homology and phylogeny of chondrichthyan tooth enameloid. Journal of Morphology. 2007 Jan;268(1):33–49.

55. Cope ED. Zittel’s manual of paleontology. American Naturalist. 1887; p.1014–1019.

56. Vogt C. Ueber die Siphonophoren. Zeitschrift für wissenschaftliche Zoologie. 1852;3:522–525.

57. Hamel MH. A new lower actinopterygian from the early Permian of the Paraná Basin, Brazil. Journal of Vertebrate Paleontology. 2005 Mar 11;25(1):19–26.

58. Štamberg S. A new aeduellid actinopterygian from the Lower Permian of the Krkonoše Piedmont Basin (Bohemian Massif) and its relationship to other Aeduellidae. Bulletin of Geosciences. 2010;85(2):183–98.

59. Chen D, Janvier P, Ahlberg PE, Blom H. Scale morphology and squamation of the Late Silurian osteichthyan Andreolepis from Gotland, Sweden. Historical Biology. 2012;24(4):411–23.

60. Ivanov AO, Nestell MK, Nestell GP. Middle Permian fish microremains from the early Capitanian of the Guadalupe Mountains, West Texas, USA. Micropaleontology. 2015 Jan 1;61(4-5):301–12.

61. Chrząstek A. Vertebrate remains from the Lower Muschelkalk of Raciborowice Górne (North-Sudetic Basin, SW Poland). Geological Quarterly. 2010 Mar 27;52(3):225–38.

62. Štamberg S. Actinopterygians of the Stephanian sediments of the Krkonoše Piedmont Basin (Bohemian Massif) and their palaeobiogeographic relationship. Bulletin of Geosciences. 2016;91(1).

63. Schultze HP. Marine to onshore vertebrates in the Lower Permian of Kansas and their paleoenvironmental implications. The University of Kansas Paleontological Contributions. 1985;113:1–18.

64. Esin DN. The scale cover of Amblypterina costata (Eichwald) and the palaeoniscid taxonomy based on isolated scales. Paleontological Journal. 1990;2(9). Russian.

65. Sasagawa I, Ishiyama M, Yokosuka H, Mikami M, Uchida T. Tooth enamel and enameloid in actinopterygian fish. Frontiers of Materials Science in China. 2009;3(2):174.

66. Young J. On the affinities of Platysomus and allied genera. Quarterly Journal of the Geological Society. 1866;22(1-2):301–17.

67. Blainville HMD. Sur les ichthyolités ou les poissons fossiles. Nouvelle Dictionaire d’Histoire naturelle, Paris. 1818;37:310–95. French.

68. Esin, D. N. 1997. Peculiarities of trophic orientation changes in palaeoniscoid assemblages from the Upper Permian of the European part of Russia. Modern Geology 21:185–196.

69. Perea D, Soto M, Veroslavsky G, Martínez S, Ubilla M. A Late Jurassic fossil assemblage in Gondwana: biostratigraphy and correlations of the Tacuarembó Formation, Parana Basin, Uruguay. Journal of South American Earth Sciences. 2009;28(2):168–79.

70. Lombardo C, Tintori A, Tona D. A new species of Sangiorgioichthys (Actinopterygii, Semionotiformes) from the Kalkschieferzone of Monte San Giorgio (Middle Triassic; Meride, Canton Ticino, Switzerland). Bollettino della Società Paleontologica Italiana. 2012;51(3):203–12.

71. Cavin L, Deesri U, Suteethorn V. The Jurassic and Cretaceous bony fish record (Actinopterygii, Dipnoi) from Thailand. Geological Society, London, Special Publications. 2009;315(1):125–39.

72. Cuny G, Cobbett AM, Meunier FJ, Benton MJ. Vertebrate microremains from the Early Cretaceous of southern Tunisia. Geobios. 2010;43(6):615–28.

73. Andreev PS. Convergence in dental histology between the Late Triassic semionotiform Sargodon tomicus (Neopterygii) and a Late Cretaceous (Turonian) pycnodontid (Neopterygii: Pycnodontiformes) species. Microscopy research and technique. 2011;74(5):464–79.

74. Shimada K, Williamson TE, Sealey PL. A new gigantic pycnodont fish from the Juana Lopez Member of the Upper Cretaceous Mancos Shale of New Mexico, USA. Journal of Vertebrate Paleontology. 2010;30(2):598–603.

75. Szabó M, Gulyás P, Ősi A. Late Cretaceous (Santonian) pycnodontid (Actinopterygii, Pycnodontidae) remains from the freshwater deposits of the Csehbánya Formation (Iharkút, Bakony Mountains, Hungary). Annales de Paléontologie. 2016;2:123–134.

76. Kriwet J, Schmitz L. New insight into the distribution and palaeobiology of the pycnodont fish Gyrodus. Acta Palaeontologica Polonica. 2005;50(1).

77. Knoll F, Cuny G, Mojon PO, López-Antoñanzas R, Huguet D. A new vertebrate-, ostracod-, and charophyte-bearing locality in the Middle Jurassic of the Grands Causses (southern France). Proceedings of the Geologists’ Association. 2013;124(3):525–9.

78. Sweetman SC, Goedert J, Martill DM. A preliminary account of the fishes of the Lower Cretaceous Wessex Formation (Wealden Group, Barremian) of the Isle of Wight, southern England. Biological journal of the Linnean Society. 2014;113(3):872–96.

79. Johnson GD. Hybodontoidei (Chondrichthyes) from the Wichita-Albany Group (Early Permian) of Texas. Journal of Vertebrate Paleontology. 1981;1(1):1–41.

80. Hodnett JP, Elliott DK, Olson TJ. A new basal hybodont (Chondrichthyes, Hybodontiformes) from the Middle Permian (Roadian) Kaibab Formation, of northern Arizona. New Mexico Museum of Natural History and Science Bulletin. 2013;60:103–8.

81. Ivanov AO, Lebedev OA. Permian chondrichthyans of the Kanin Peninsula, Russia. Paleontological Journal. 2014 Dec 1;48(9):1030–43.

82. Solodukho MG. Nakhodki predstaviteley sem. Platysomidae v verkhnekazanskikh otlozheniyakh okrestnostey d. Pechishchi (Tatarskaya ASSR). Uchyonnye Zapisi Kazanskogo Gosudarstvennogo Universiteta im. V.I. Ul’yanova Lenina. 1951;111:157–159. Russian.

83. Westoll TS. The Permian fishes *Dorypterus* and *Lekanichthys*. Proceedings of the Zoological Society of London, Series B. 1941;111:39–58.

84. Cione AL, Gouiric-Cavalli S, Mennucci JA, Cabrera DA, Freije RH. First vertebrate body remains from the Permian of Argentina (Elasmobranchii and Actinopterygii). Proceedings of the Geologists’ Association. 2010 Jan 1;121(3):301–12.

85. Schindler, T. *‘Elonichthys’* palatines n. sp., a new species of actinopterygians from the Lower Permian of the Saar-Nahe Basin (SW-Ger many). New Research on Permo-Carboniferous Faunas. Pollichia-Buch. 1993;29:67–81.

86. Zajíc J. Vertebrate biozonation of the Permo-Carboniferous lakes of the Czech Republic – new data. Acta Musei Reginaehradecensis S.A. 2004;30:15–16.

87. Zajíc, J. 2007. Carboniferous Fauna of the Krkonoše Piedmont Basin. Acta Musei Reginaehradecensis S.A. 2007;32:11–6.

88. Štamberg S. Discovery of skeletal fragments of a large amphibian and other Permian fauna from locality Arnultovice in the Krkonoše Piedmont Basin. Geoscience Research Reports. 2014;47:94–7.

89. Heintz A. Fischreste aus dem Unterperm Norwegens. Norsk geologisk tidsskrift. 1934;14(1-2):176–94.

90. Magaritz M. A new explanation for cyclic deposition in marine evaporite basins: meteoric water input. Chemical geology. 1987;62(3-4):239–50.

91. Peryt TM, Scholle PA. Regional setting and role of meteoric water in dolomite formation and diagenesis in an evaporite basin: studies in the Zechstein (Permian) deposits of Poland. Sedimentology. 1996;43(6):1005–23.

92. Kendall AC. Evaporites. In: Walker, R. G. (Ed.), Facies Models, 2nd edn. Geoscience Canada. 1984; 259–296 p.

93. Weissflog L, Elansky NF, Kotte K, Keppler F, Pfennigsdorff A, Lange CA, Putz E, Lisitsyna LV. Late permian changes in conditions of the atmosphere and environments caused by halogenated gases. InDoklady Earth Sciences. 2009;425:291–295.

